# Anti-biofilm activity of new low molecular weight compounds produced by *Lactiplantibacillus plantarum* SJ33 against *Klebsiella pneumoniae*

**DOI:** 10.1101/2024.12.03.626605

**Authors:** Amrita Ray Mohapatra, Divya Lakshmanan, Kadirvelu Jeevaratnam

**Affiliations:** Dept. of Biochemistry and Molecular Biology, School of Life Sciences, Pondicherry University, R. V. Nagar, Kalapet, Puducherry – 605014

**Author notes:** **Corresponding Author** Prof. K. Jeevaratnam.

**Keywords:** C1 and C2, Gram-negative bacteria, Biofilm, Anti-biofilm, multi-drug resistance, Scanning electron microscopy

## Abstract

Community-acquired infections are partly attributed to biofilms formed by drug resistant bacteria signifying major health problems. Hence, there is a critical need for the development of new antibiotics with antibacterial activities. In this study, we investigated the biofilm inhibiting characteristics of antibacterial compounds C1 and C2 confirmed as (3-amino-5-hydroxy-6-(hydroxymethyl)-4-(1-hydroxyprop-2-yn-1-yl)-3,3a,4,5,6,7a-hexahydro-7H-indazol-7-one) and (1-(dimethylamino)-3-hydroxy-3-((2-hydroxypropan-2-yl)oxy)-1 (methylamino)butan-2-one) which were previously characterized and identified by our group. This compound exhibits antibacterial activity against Gram-negative bacteria including activity against clinical isolates of *Klebsiella pnemoniae* and *Escherichia coli*. Bactericidal activity against *K. pnemoniae* was revealed by scanning electron microscopic (SEM) images that showed cellular leakage by C1 and C2 which is probably the major cause of its antibacterial activity. These findings were supported by antibacterial assays of susceptible as well as resistant pathogens. Biofilm assays also proved anti-biofilm activity of C1 and C2 on biofilm inhibition as well as dispersion of mature biofilms using crystal violet assay and the results were confirmed by fluorescence and SEM observation. In conclusion, our results showed that C1 and C2 exert bactericidal action on target cells by membrane disruption. Biofilm inhibition and dispersion of preformed biofilms was also revealed by microscopic images, making it an interesting alternative for commercial antibiotics for the development of new antibacterial therapies. The bioactive compounds C1 and C2 not only inhibit *K. pnemoniae* growth but also eradicate the biofilm formation thus reducing the virulence of *K. pnemoniae* pathogen.

## INTRODUCTION

The inappropriate and abandoned use of commercial antibiotics in medicine resulted in the occurrence of multi-drug resistant (MDR) strains, which have become a worldwide health concern. MDR strains are those pathogens that obtained resistant to one or more classes of antibiotics. MDR pathogens are limiting the effectiveness of many antibiotics and are involved with several nosocomial and community-acquired infections (Iseppi *et al*., 2020).

Biofilm formation is considered as a major factor in development of infections and one of the resistance strategies of many pathogenic bacteria which makes them more difficult to treat than the planktonic bacteria (Zamani *et al*., 2017). Biofilm is an intricate polymeric matrix of microbial communities composed of polysaccharides, proteins and other components in which cells bind together strongly and also enable microbes to adhere on to surfaces that can tolerate hostile conditions such as antimicrobial agents and host defenses. Therefore, biofilm formation is one of the indirect mechanism by which microorganisms attains resistant to antibiotics where they also transfer resistance genes within the biofilm micro-community (Famuyide *et al*., 2019a). Many Gram-negative bacteria of genera such as *Escherichia*, *Klebsiella*, *Pseudomonas* and *Salmonella* could form biofilm that cause serious infections. Nearly, two-third of bacterial infections in humans and majority of microbial infections are caused by the biofilms (Famuyide *et al*., 2019b). The resistant nature of biofilm is a matter of immense concern for medicine and health care industry and many studies have been carried out to explore novel and effective anti-biofilm agents (Zamani *et al*., 2017).

Gram-negative pathogens such as *Klebsiella pneumoniae*, *Escherichia coli* and *Pseudomonas aeruginosa* are among the leading causes of nosocomial infections throughout the world and have developed the resistance to antibiotics. *P. aeruginosa* and *K. pneumoniae* are the causes of several potentially fatal infections, including pneumonia, sepsis, wound, surgical site infections and meningitis. *K. pneumoniae* and *E. coli* pathogens have been reported to attain resistance to most of the generally used traditional antibiotics (Yarlagadda *et al*., 2018). The presence of the outer membrane in Gram-negative bacteria offers further difficulty towards identification of novel antibacterial compounds and have abilities to develop new mechanisms of resistance (Choi, Britigan and Narayanasamy, 2019). The increase of antimicrobial resistance to antibiotics is a significant health problem, hence there is an urgent need to develop new antibacterial compounds as alternative therapeutic agents that can block resistance to eliminate these resistant strains and improve treatment of infections (Haroun and Al-Kayali, 2016).

Low-molecular-weight (LMW) antimicrobial compounds are known to be produced by most of the lactic acid bacteria (LAB) strains which are generally considered as safe (Jeevaratnam *et al*., 2015). Few *Lactobacillus spp.* produce antimicrobial compounds that not only inhibit bacterial growth but also has inhibitory effect on biofilm formation by *K. pneumoniae* (Yudani *et al*., 2019a). While few other LMW antimicrobial compounds produced by *Lactobacillus* spp. have not been characterized and analyzed extensively due to their less concentration and low yield. Furthermore, only limited studies have been performed on antimicrobial compounds produced by *Lactobacillus* spp. to evaluate their potential as natural therapeutic agents to overcome microbial drug resistance in clinical infections (Yudani *et al*., 2019b).

Previously, we purified and identified the structures of two novel antibacterial compounds C1 and C2 (Figure 1) with antibacterial and anti-biofilm activity against susceptible *S. aureus* (MSSA) and MRSA. Additionally, we also demonstrated that the LMW compound can be used as antibacterial coating on urinary catheters, inhibiting bacterial colonization on medical devices (Mohapatra et al., manuscript communicated). In the present work, we further studied the antibacterial characteristics, including anti-biofilm activity of novel LMW against Gram-negative pathogens i.e. *K. pneumoniae* and *E. coli*. This study aimed to develop a strategy to address the emergence of antibiotic resistance through bactericidal and anti-biofilm properties of compounds for treating biofilm-associated infections as well as reduce bacterial pathogenesis.

**Figure 1.**
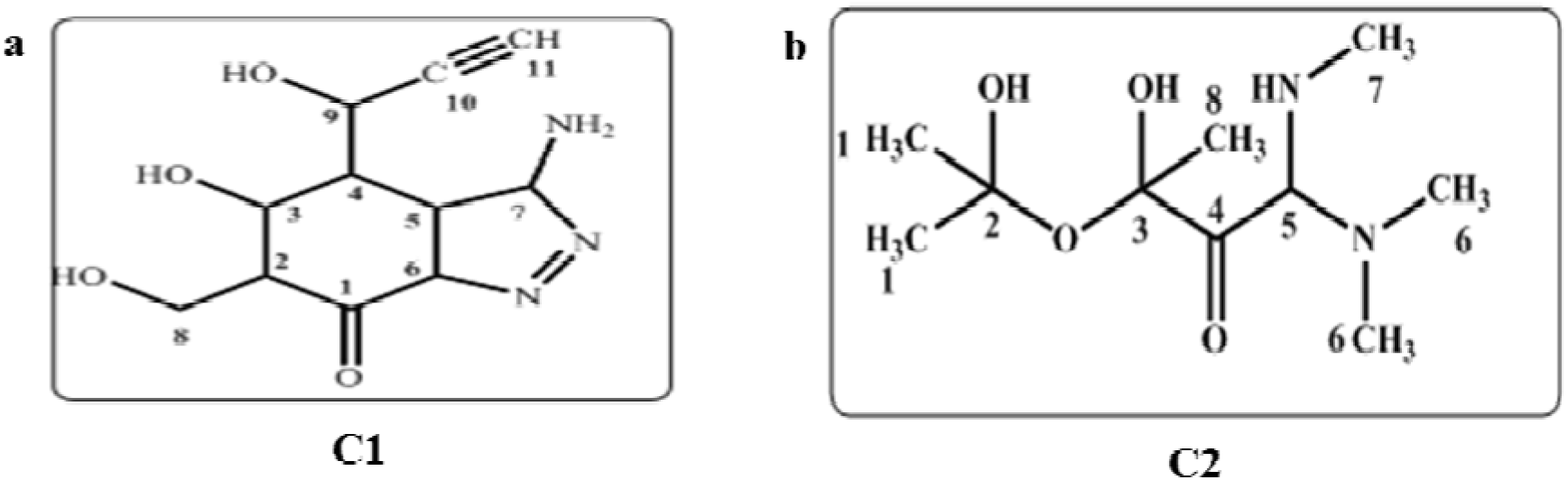
Structure of LMW compounds (a) **C1** predicted as **3-amino-5-hydroxy-6-(hydroxymethyl)-4-(1-hydroxyprop-2-yn-1-yl)-3,3a,4,5,6,7a-hexahydro-7H-indazol-7-one** and (b) **C2** identified as **1-(dimethylamino)-3-hydroxy-3-((2-hydroxypropan-2-yl)oxy)-1 (methylamino)butan-2-one**.

## MATERIAL AND METHODS

### Bacterial Strains and growth conditions

Multi-resistant clinical isolates of *K. pneumoniae* and *E. coli* were collected from Mahatma Gandhi Medical College and Research Institute (MGMCRI), Pondicherry, India. The clinical isolates were grown in Mueller-Hinton broth (MHB) (HiMedia, India). *K. pneumoniae* (MTCC 109), *K. pneumoniae* (MTCC 3384) and *E. coli* (MTCC 728) susceptible bacterial strains used in this study were obtained from Microbial Type Culture Collection (MTCC), Institute of Microbial Technology, Chandigarh, India. All pathogenic strains were cultured in Mueller-Hinton broth (MHB) and agar (MHA). The producer strain, *Lactobacillus plantarum* SJ33 was grown in Man Rogosa Sharpe (MRS) broth and maintained in 50% glycerol at −80 °C.

### Structure of LMW compounds

Purified antibacterial compounds obtained from reverse phase HPLC were structurally characterized by spectroscopic studies which involve 1D NMR and 2D NMR (COSY, NOESY, HSQC), Infrared (FT IR) spectroscopy and molecular masses of C1 and C2 were confirmed by HR ESI-MS analysis in our previous study (Ray Mohapatra *et al*., 2021).

### Antibacterial assay

Antibacterial activity of fractions obtained after purification was tested against both sensitive and drug-resistant Gram-negative pathogens. Antibacterial efficacy of two purified compounds C1 and C2 were screened at a single concentration against bacterial strains and clinical isolates using agar well diffusion assay (Parente *et al*., 1995). The bacterial cultures were grown overnight in broth media (1% v/v) inoculated in agar (1.2% MHA). The agar media inoculated with pathogens were poured into Petri plates and 6 mm diameter wells were bored into the plates. The Petri plates were incubated for overnight at 37°C and the diameters of the inhibition zone was measured.

### Determination of minimum inhibitory concentrations

The Minimum inhibitory concentration (MIC) of compounds C1 and C2 were determined by broth micro-dilution method (Zgoda and Porter, 2001; Jiang *et al*., 2013). Compounds were serially diluted to obtain different concentrations, dilutions were distributed into the wells of the 96 well microtiter plate and then 100 µL of bacteria (10^6^ CFU/mL) was added to each well to attain the desired final concentrations in range 0.37 to 24 mM. The plate was incubated at 37°C for 24 h and cell density was measured at 600 nm. Ciprofloxacin (Cip) with similar concentrations as for the antibacterial compounds was used as an antibiotic standard. The MIC value was evaluated as the lowest concentration at which the compounds totally inhibit the bacterial growth (Valgas *et al*., 2007). All experiments were performed in triplicates in three independent experiments.

### Mechanism of action

The overnight grown culture of *K. pneumonia* MTCC 109 and *K. pneumoniae* clinical isolate were treated with C1 and C2 compounds at MIC and incubated for 6 h at 37°C to analyze the morphological changes under SEM. The cells were harvested by centrifugation at 8,000 x *g* for 5 mins, the cell pellet was washed twice with phosphate buffer saline (PBS) and fixed with 2.5% (v/v) glutaraldehyde. After fixation the cells were centrifuged (8,000 x *g* for 5 mins) and washed with PBS, then the cells were treated with 1% osmium tetroxide for 1 h at room temperature, the centrifugation and washing step was repeated again. Finally the cells were dehydrated through a gradient series of ethanol (50%, 75%, 90%, 95% and 100%) which were dried to completely remove the supernatant. The cells were lyophilized and coated to be examined under HR SEM (FEI Quanta FEG 200, USA) (Kim *et al*., 2009).

### Biofilm assays

#### Congo red assay

Congo red agar (CRA) medium was prepared with brain heart infusion broth 37 g/L, sucrose 50 g/L, agar 10 g/L and Congo Red indicator 8 g/L supplemented with 5% sucrose and congo red stain was sterilized (121°C for 15 minutes) separately from the other medium constituents. Then it was added to the autoclaved brain heart infusion agar with sucrose at 55°C. CRA plates were inoculated in streaks with test organisms and incubated at 37°C for 24 h aerobically (Mathur *et al*., 2006; Hassan *et al*., 2011). The experiment was performed independently in triplicates.

#### Biofilm formation

Biofilm formation of the bacterial pathogenic strains was quantified by crystal violet assay in 96-well microtitre plates. The diluted (1:100) overnight cultures of pathogenic strains (∼10^6^ CFU/mL) were added to microtitre plates and kept for static incubation for 24 h at 37°C. After incubation, plates were washed gently with PBS to remove the non-adhered cells, then stained with 0.1% (w/v) crystal violet for 10 min and washed with PBS. The bound stain was dissolved in 33% (v/v) acetic acid and measured using microplate reader at 590 nm (Versa max microplate reader) to detect the density of biofilms formed (Tong *et al*., 2014).

#### Biofilm inhibition by antibacterial compounds

The inhibitory effect of two bioactive compounds C1 and C2 were examined at sub-MIC (1.5 mM) on biofilm formed by strong biofilm formers. The disruption of biofims formed by susceptible as well as resistant clinical isolates was evaluated quantitatively in a similar way by crystal violet staining as mentioned above. Strong biofilm formers including *K. pneumoniae* MTCC 109, *E. coli* MTCC 728 and clinical isolates of *K. pneumonia* and *E. coli* were used in this study for analyzing the anti-biofilm activity of the antibacterial compounds C1 and C2. The assay was carried out in triplicates in three different experiments.

#### Dispersion of established preformed biofilms

Biofilms were allowed to be formed for 24 h prior to addition of compounds C1 and C2. Biofilm formation was achieved by transferring 100 μL of bacterial culture (prepared as described above) into the wells of polystyrene microtitre plates in triplicates. The microtitre plates were incubated for 24 h at 37°C to allow cell attachment and initiate biofilm formation. Following incubation, the plate was washed with sterile distilled water to remove non-adhered cells and 100 μL of C1 and C2 solution at sub-MIC (3mM) was added to each well. Equal volumes of sterile distilled water were added as negative control. After the treatment of preformed biofilms, the plates were incubated for overnight. Following incubation, the biofilms were assessed using the crystal violet assay.

#### Visualization of biofilms under Fluorescence microscope and SEM

The overnight cultures of biofilm forming *K. pneumoniae* MTCC 109 strain were diluted (1:100) with MHB and added to 12-well plate with and without C1 and C2 at sub-MIC and the plates were incubated for 24 h at 37°C. The biofilms after incubation were viewed under fluorescence microscope with Live/Dead staining (EtBr/AO) (Tong *et al*., 2014). The disruptions of preformed biofilms were observed after 24 h biofilm development and then addition of the compounds.

The biofilms to be examined under SEM were prepared on coverslips, the coverslips were incubated in 12-well plate with the samples untreated and treated with sub-MIC concentration in a similar way for 24 h at 37°C. After incubation the coverslips were washed with PBS, dried and fixed with glutraldehyde (2.5%). The fixed coverslips were finally dehydrated with gradient series of ethanol (50%, 75%, 90%, 95% and 100%) to visualize the formation and inhibition of biofilm in *K. pneumoniae* MTCC 109 strain under HR SEM (Lakshmanan *et al*., 2019).

## RESULTS

### Structure of antibacterial compounds

The purified LMW compounds produced by *Lactobacillus plantarum* SJ33 strain were purified by RP HPLC and were structurally identified (Figure 1A and B) by NMR, IR and HR ESI-MS data in our previous study (Mohapatra et.al. manuscript communicated). The molecular structure of the bioactive compound C1 was previously elucidated as **3-amino-5-hydroxy-6-(hydroxymethyl)-4-(1-hydroxyprop-2-yn-1-yl)-3,3a,4,5,6,7a-hexahydro-7H-indazol-7-one** and the molecular formula was C_11_H_15_N_3_O_4_. The structure of antibacterial compound C2 was confirmed as **1-(dimethylamino)-3-hydroxy-3-((2-hydroxypropan-2-yl)oxy)-1 (methylamino)butan-2-one** and the molecular formula of C2 was C_10_H_23_N_2_O_4_ (Figure 1) reported in our earlier study (Ray Mohapatra *et al*., 2021).

### Antibacterial assay

The activity of compounds produced by *Lactobacillus plantarum* SJ33 were determined against susceptible as well as resistant Gram-negative bacteria and showed good antibacterial activity against *K. pneumoniae* and *E. coli* (Table 1). Table 1 clearly showed C2 was more effective against susceptible *K. pneumoniae* and *E. coli* strains whereas C1 revealed better antibacterial activity against clinical isolates of *K. pneumoniae* and *E. coli* at equal concentration. The supplementary figure also showed potent activity of C1 against resistant *K. pneumoniae* and *E. coli* clinical isolate (Figure S1).

### Determination of minimum inhibitory concentration

The inhibitory potential of compounds C1 and C2 were determined against both susceptible and resistant Gram-negative pathogens. C1 and C2 were found to be effective against *K. pneumoniae* MTCC 109 and clinical isolate of *K. pneumoniae* that showed more than 90% of inhibition at 12 mM (Figure 2). The susceptible pathogenic strains were found to be more sensitive towards C2 while C1 was effective against resistant bacteria isolates at MIC (Figure 2). Ciprofloxacin, the standard antibiotic was also effective at lower concentrations as compared to the compounds C1 and C2.

**Figure 2.**
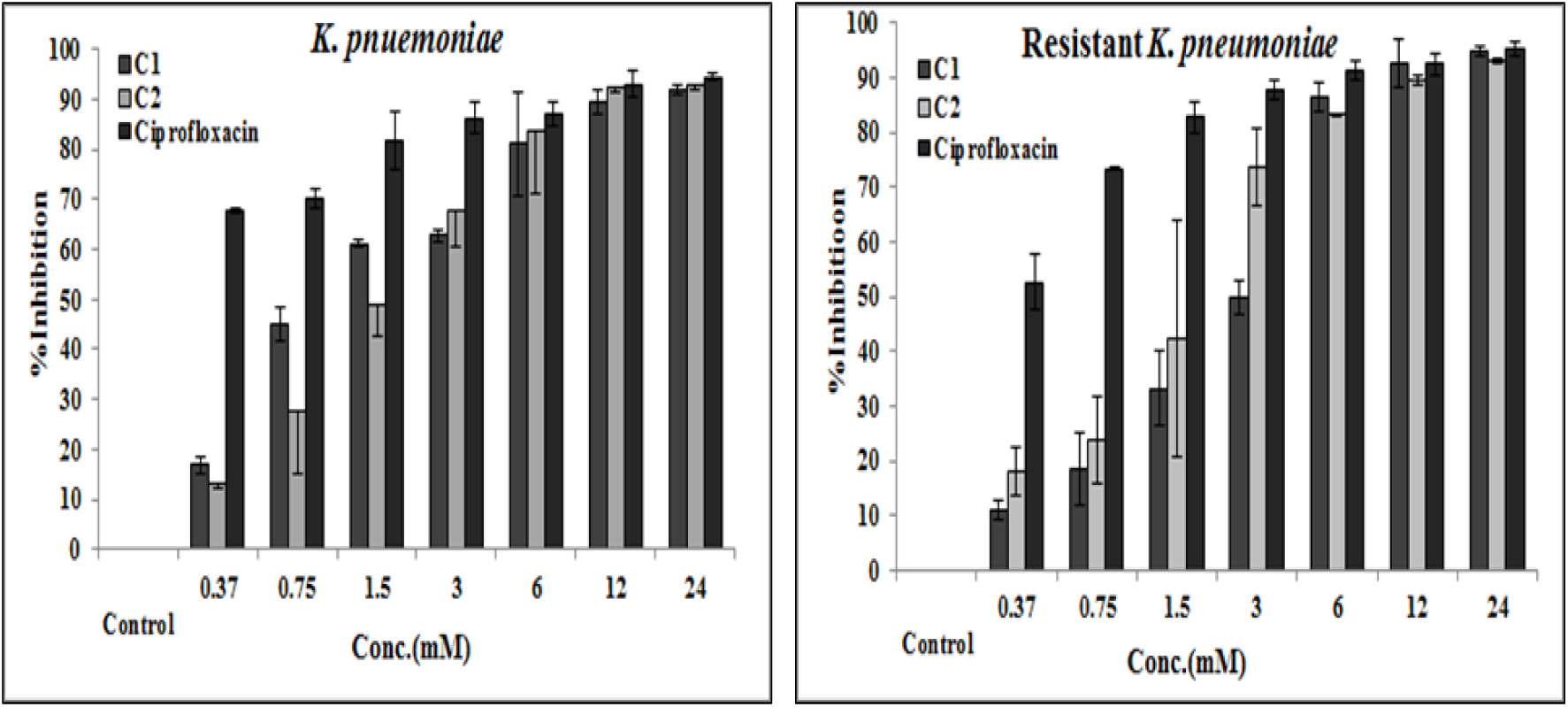
Minimum inhibitory concentration of compounds C1 and C2 against *K. pneumoniae* MTCC 109 and drug-resistant *K. pneumoniae*.

### Mechanism of action

The activity of C1 and C2 on *K. pneumoniae* MTCC 109 cells were viewed under SEM and deformation of cell morphology was clearly visible (Figure 3B and 3C) after the treatment of antibacterial compounds at MIC (12 mM). After incubation with C1 and C2 for 6 h the membrane integrity was lost, membrane disruption and damage was evidently visible on the surface of the target pathogen (Figures 3B and 3C). Figure 3B and 3C clearly showed the leakage of inner contents in compound treated *K. pneumoniae* MTCC 109 cells when compared to the untreated cells in Figures 3A.

**Figure 3.**
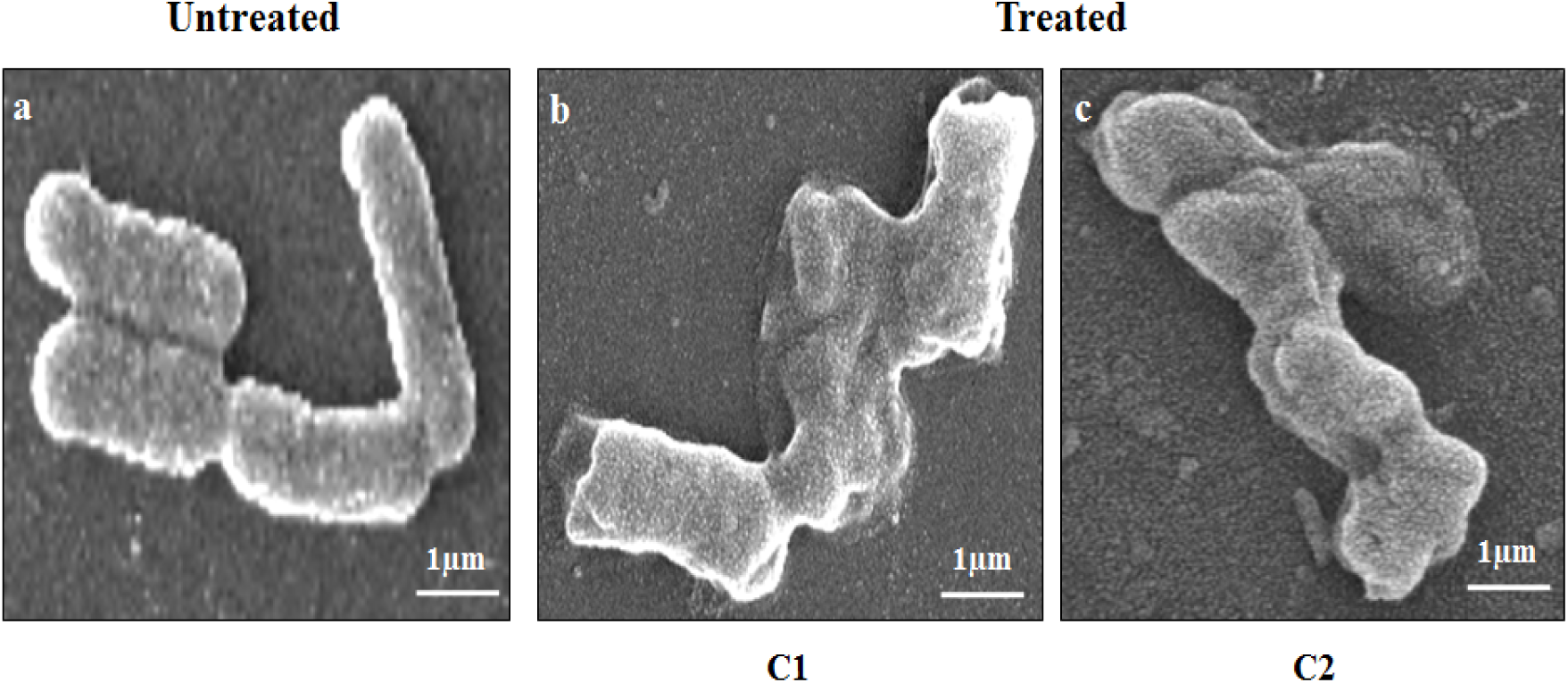
Bactericidal action of compounds C1 and C2 against *K. pneumoniae* MTCC 109 revealed by SEM analysis. (A) Untreated cells. Cells treated with (B) C1 and (C) C2 at 3mM concentration.

### Biofilm assays

#### Congo red assay

Congo red agar (CRA) method showed streaked bacterial pathogens that displayed black colonies on Congo red agar plates as distinctive biofilm producing strains, but no dry crystalline morphology was observed. However, some bacterial strains such as *P. aeruginosa* MTCC 2295 showed red or pink colonies (Figure S2). Only few isolates displayed red (pink or orange) colonies after incubation at 37°C for 24 to 48 h.

#### Biofilm inhibition by LMW compounds

The anti-biofilm activity of C1 and C2 was explored against biofilm forming Gram-negative pathogens. The activity of purified compounds C1 and C2 at sub-MIC on biofilm formation and on established mature biofilms was quantified by crystal violet assay. The results showed biofilm inhibition at 3 mM and biofilm dispersion at 6 mM of *K. pneumoniae* MTCC 109 and *K. pneumoniae* clinical isolate. The percentage of biofilm inhibition by C1 was found to be in a range of 20 to 45% while C2 reduced the biofilm formation by 30 to 60% at 3 mM (Figure 4). Notable, biofilm inhibition and biofilm dispersion was found against susceptible as well as clinical isolate of *K. pneumoniae* specifically, against *K. pneumoniae* MTCC 109 strain i.e. nearly 60%, whereas the lowest percentage of biofilm inhibition and dispersion was seen against *E. coli* clinical isolate (Figure 4A and 4B). The quantitative results of crystal violet staining clearly showed that C2 have better biofilm inhibition potential than C1, while C1 efficiently dispersed preformed mature biofilms as compared to C2 (Figure 4B).

**Figure 4.**
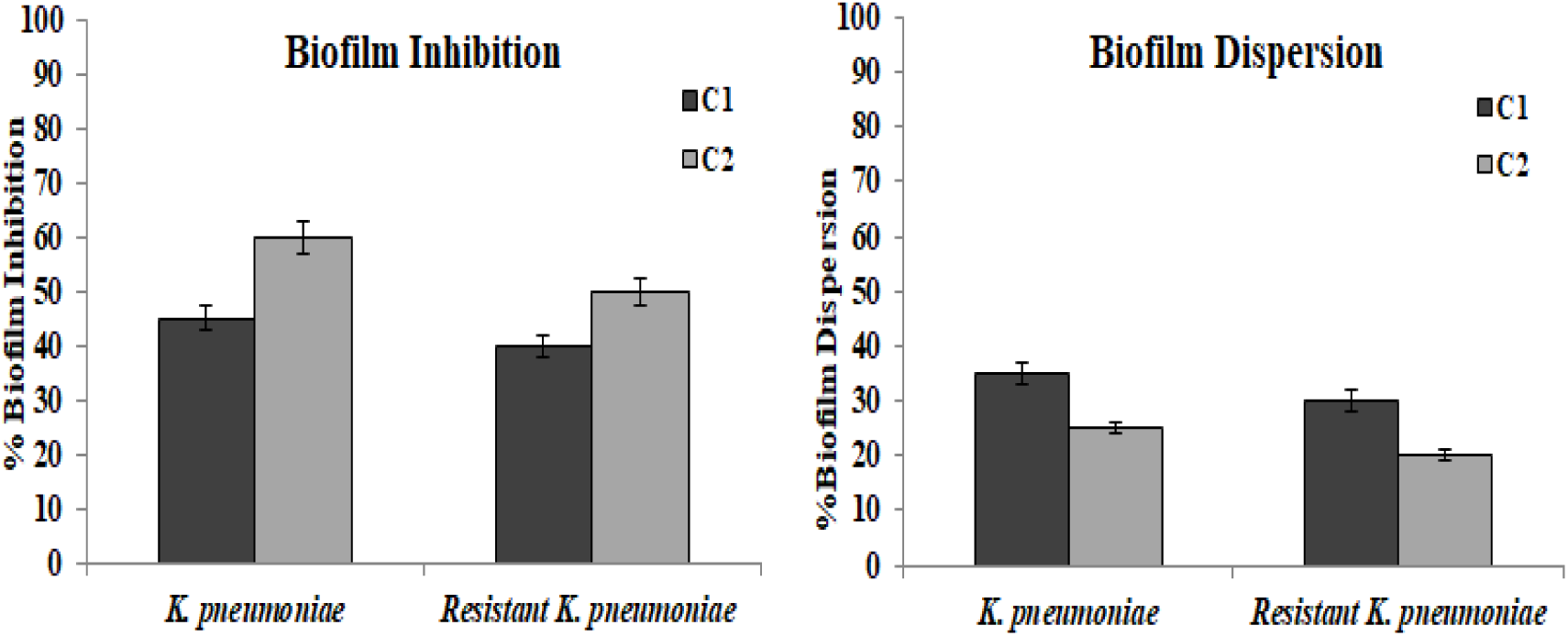
Antibiofilm activity of compounds C1 and C2 against Gram-negative pathogens using crystal violet assay. (A) Biofilm inhibition and (B) dispersion of preformed biofilms.

#### Visualization of biofilms under Fluorescence microscope and SEM

The results of crystal violet assay were further confirmed by fluorescence and SEM images which demonstrated both inhibition as well as dispersion of preformed biofilms of *K. pneumoniae* MTCC 109 strain (Figure 5). Biofilms formed by *K. pneumoniae* clinical isolate also showed inhibition and dispersion by compounds C1 and C2 (Figure 5). SEM images also clearly demonstrated inhibition of biofilms formed by *K. pneumoniae* MTCC 109 strain (Figure 6) which confirmed the anti-biofilm activity of both compounds against the biofilm forming Gram-negative pathogen.

**Figure 5.**
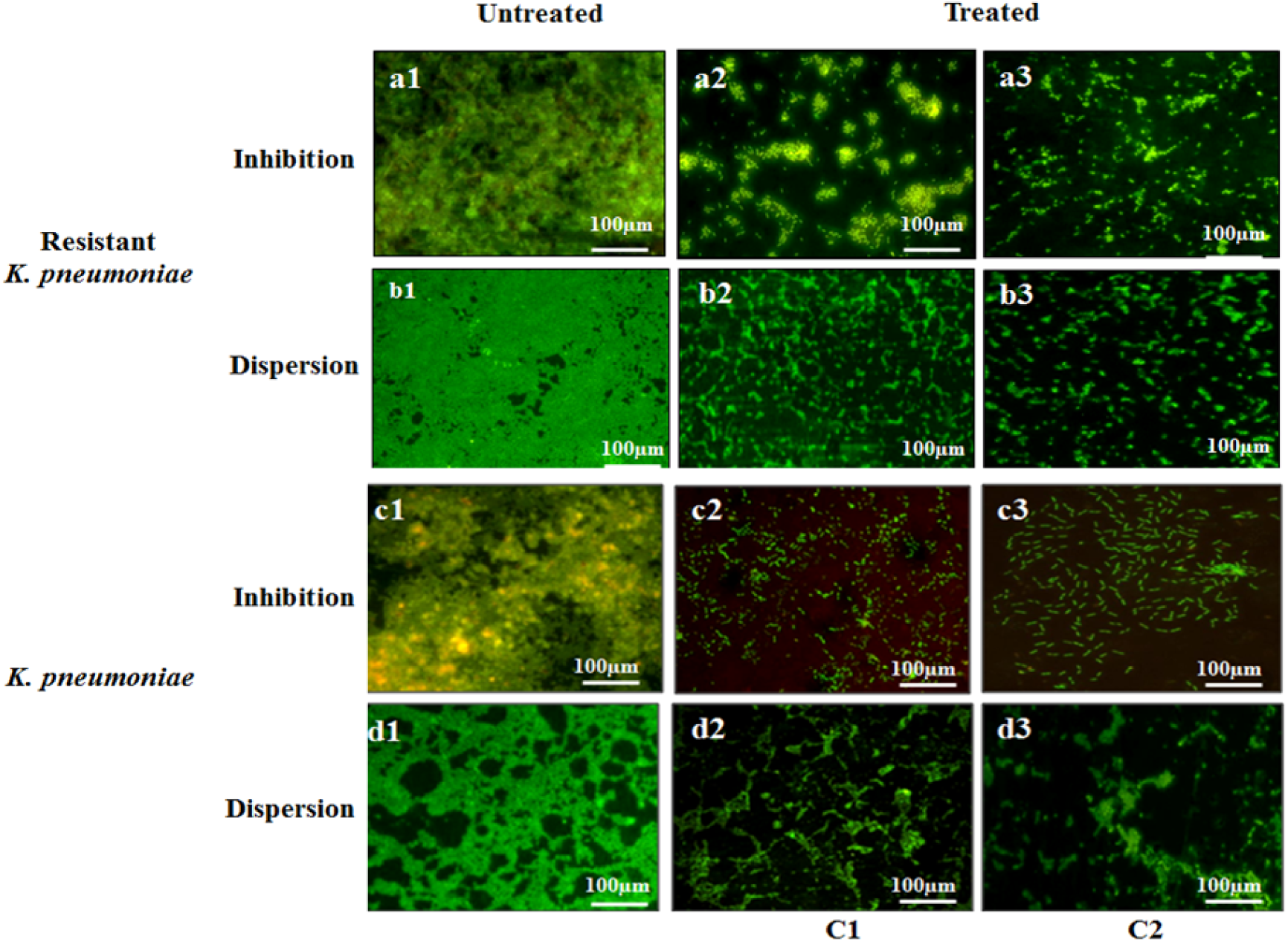
Antibiofilm efficacy of compounds C1 and C2 against *K. pneumoniae* MTCC 109 and drug-resistant *K. pneumoniae* viewed under Fluorescence microscope. Inhibition and dispersion of susceptible and resistant *K. pneumoniae* biofilms and preformed 24 h old biofilms grown either in absence (control) or presence (treated) of C1 and C2 at sub-MIC.

**Figure 6.**
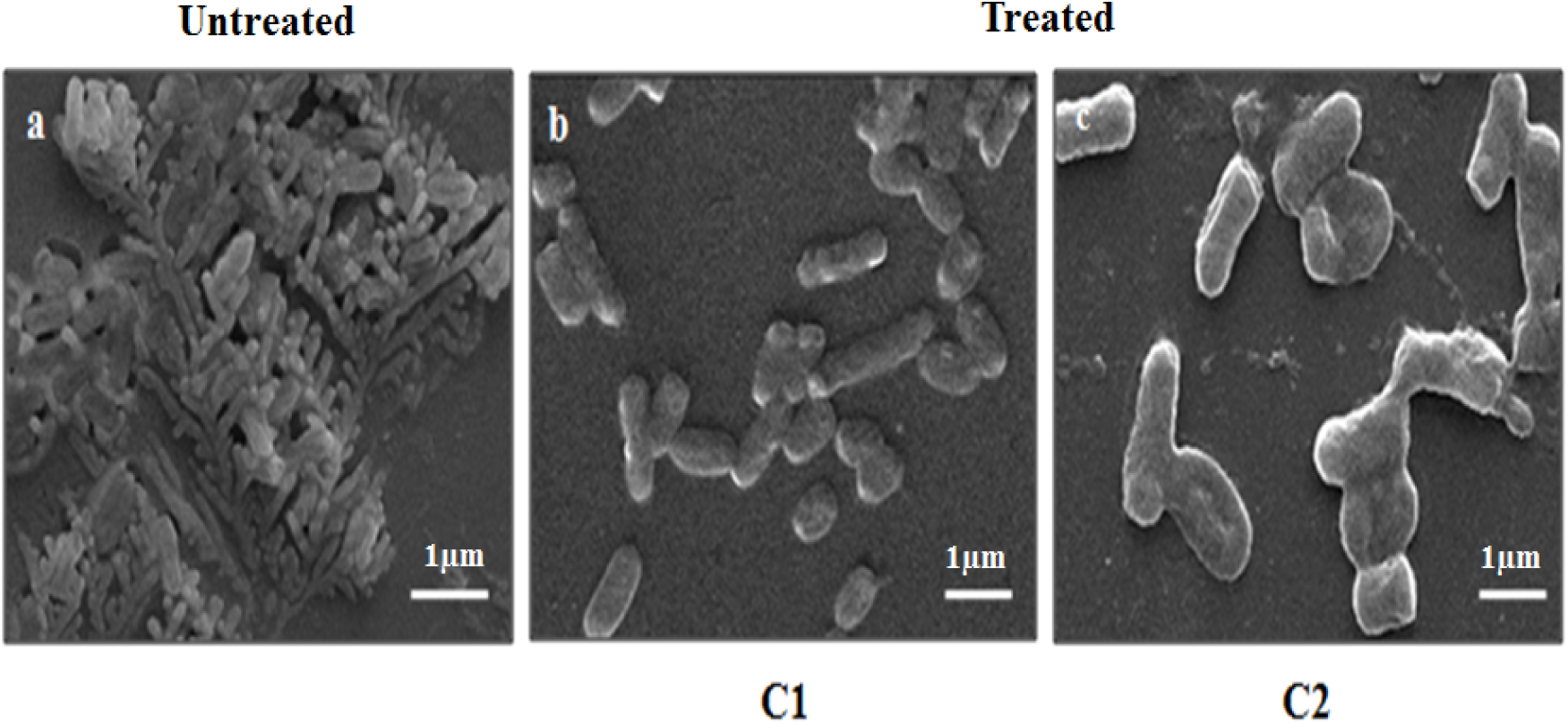
Microscopic analysis of biofilm inhibition against *K. pneumoniae* MTCC 109 strain by compounds C1 and C2. SEM images showed (A) biofilm formation and (B) C1 and (C) C2 treated *K. pneumoniae* showed reduction or disruption of biofilms.

## DISCUSSION

Gram-negative bacteria are among the pathogens that cause hospital-acquired and clinical infections, which have emerged as a major threat to healthcare sector. Additionally, *K. pneumoniae* causes nosocomial infections and has also appeared as one of the means of community-acquired infections. Antimicrobial agents that are effective against multi-drug-resistant pathogens are limited (Haroun and Al-Kayali, 2016). Thus, it is important to search and develop novel compounds that potentiate activity of antibiotics on these drug-resistant pathogens. Therefore, combating multi-drug resistance which selectively interferes with mechanism that regulate virulence are difficult to treat (Lesic *et al*., 2007) and urgently require alternative mechanistic approach that can combat antibiotic resistance to the improve chemotherapeutic methods.

The current study showed antibacterial activity of two recently identified low molecular weight compounds (Mohapatra et al., manuscript communicated) against both susceptible as well as drug resistant Gram-negative bacteria. However, there were various antimicrobial compounds produced by different LAB strains which were previously reported i.e. Reuterin produced by LAB strains such as *Lactobacillus brevis*, *Lactobacillus buchneri*, *Lactobacillus collinoides*, *Lactobacillus coryniformis* and was known as well characterized low molecular antimicrobial compound till date (Talarico *et al*., 1988; Talarico and Dobrogosz, 1989; Vollenweider *et al*., 2003; Cleusix *et al*., 2007; Jeevaratnam *et al*., 2015). The compounds characterized in this study demonstrated antibacterial effect against Gram-negative bacterial strains. Similar activities were previously reported for other antimicrobial substances isolated from different LAB sources that showed activity against Gram-negative pathogens (Bernet-Camard *et al*., 1997; Yang *et al*., 1997; Tenea *et al*., 2019). Another study demonstrated cell-free supernatant of different *Lactobacillus* and *Bifidobacterium* spp. that were able to reduce the growth of drug resistant *E. coli* isolates when examined using well diffusion method (Abdelhamid, Esaam and Hazaa, 2018). In contrast to our results previous studies showed novel compound, 135C was mostly active against Gram-positive bacteria and observed poor activity against Gram-negative bacteria. The lack of activity against Gram-negative bacteria may be due to several mechanisms including outer membrane impermeability (Man *et al*., 2018). In addition, anti-parasitic agent i.e. auranofin acquires significant antibacterial activity against important Gram-positive pathogens, including MRSA but lacks antibacterial efficacy against Gram-negative bacteria as indicated by an earlier study (Thangamani, Mohammad and Abushahba, 2016).

The effect of compounds C1 and C2 on the pathogenic strains showed that the inhibitory concentration varies depending on different strains and means of determination. Comparison of MIC between C1 and C2 as well as among different bacterial pathogens revealed that the compounds were effective against both sensitive as well as drug resistant *K. pneumoniae* which was also seen in case of the standard antibiotic used in this study. Previous studies also suggested higher MIC values of antibacterial metabolites produced by *Lactobacillus* spp., specifically MIC of low molecular antimicrobial compounds produced by different LAB strains can be as high as 200-500 mM and the inhibitory concentration ranges from 8 to 500 mM depending on the target organism (Schwenninger *et al*., 2008). Reuterin produced by *L. reuteri*, a low molecular weight antimicrobial compound also showed higher active concentrations in a range of 1.4 M to 0.030 M (Spinler *et al*., 2008). Antimicrobial activities of biosurfactancts derived from *Lactobacillus* spp. against *E.coli* and drug-resistant *K. pneumoniae* demonstrated inhibitory concentration ranging between 12.5 mg/mL to 50 mg/mL (Sambanthamoorthy *et al*., 2014a) which also suggested higher MIC values. The cell free extract (CFS) of *L. plantarum* was found to inhibit diverse pathogens at concentrations ranging higher than 1750 µg/mL and 3350 µg/mL (Lin and Pan, 2019). These reports supported our present study which also showed similar results.

Besides, some earlier studies also showed MIC of plant extracts against multi-drug resistant *S. aureus* and *K. pneumoniae* strains vary from 6.25 to 50 mg/mL and 12.5 to 100 mg/mL respectively, which indicate moderate antibacterial activity and showed much higher MIC values than our results. The antibacterial action of C1 and C2 was investigated to analyze the bactericidal effects of the compounds on *K. pneumoniae* cells. The results of SEM clearly revealed membrane disruption and leakage of cellular components which confirmed the bactericidal nature of both compounds against *K. pneumoniae*. Previous study also demonstrated microbicidal activity of biosurfactants isolated from *Lactobacillus* spp. that targeted bacterial cell membrane as revealed by electron microscopic analysis (Sambanthamoorthy *et al*., 2014b). Antibacterial compounds from different bacterial strains, specifically from LAB can damage the target cells by various mechanisms but the exact mechanism for bactericidal activity is still unknown (Liu *et al*., 2017; Man *et al*., 2018).

Biofilm related infections are among major concerns than the infections caused by their planktonic counter parts. Here, we evaluated biofilm forming ability of Gram-negative pathogens and clinical isolate of *K. pneumoniae.* Recent studies suggested that biofilm formation may be an important virulence factor of *K. pneumoniae* pathogen and have the ability to form strong biofilms on surfaces (Derakhshan, Navidinia and Ahmadi, 2018). Congo red assay used for screening biofilm forming strains was simple to perform and the results were commonly based on color of colonies produced on CRA plates, which ranges from red (or pink) for non-biofilm formers to black for biofilm formers (Kaiser *et al*., 2013). This study has also examined the antibiofilm activity of reported compounds C1 and C2 against Gram-negative pathogens for biofilm inhibition as well as dispersion of 24 h old biofilms by quantitative crystal violet assay. Both the antibacterial compounds effectively inhibit and disperse *K. pneumoniae* biofilms at sub-MIC which was further confirmed using fluorescent microscopy and SEM images. These results suggested that C1 and C2 retained its anti-biofilm activity even at lower concentrations than MIC. Previous study of ASK2, a bioactive compound obtained from *Streptomyces* sp. prevented *K. pneumoniae* biofilm formation at sub-MIC, signifying that ASK2 does not target virulence gene expression (Lalitha, Raman and Rathore, 2017). Biosurfactants obtained from *Lactobacillus* spp. showed anti-biofilm property and also disrupted preformed biofilms of clinically significant pathogens as reported earlier (Sambanthamoorthy *et al*., 2014b). Previously, cell free preparations of *L. plantarum*, *L. acidophilus*, *L. helveticus* and various other LAB strains also recorded anti-biofilm activity and reduction ability against biofilms of *K. pneumoniae* and *E. coli* (Allam, 2017; Yudani *et al*., 2019b). On the other hand, the anti-biofilm activity of antibiotics ranged from 512 to 5120 mg/L, whereas some antibiotics showed anti-biofilm activity at 80 mg/L. These previous reports suggested that few antibiotics restrained biofilm formation against pathogenic strains at high range which clearly demonstrated our studied compounds to be more efficient than few conventional antibiotics. The CFS of *Lactobacillus acidophilus* also showed better antibacterial and anti-biofilm activities against *K*. *pneumoniae* pathogen than other Gram-negative bacteria as suggested by the previous study (Mohamed A El-Mokhtar *et al*., 2020).

Moreover, only few bioactive compounds produced by LAB strains demonstrated anti-biofilm activity as reported earlier. Most of the studies suggested combination of antibiotics at sub-minimal concentration to have synergistic effect on biofilms (Rachid *et al*., 2000). However, combination therapies of antibiotics and antimicrobial metabolites can help in reducing the high dosage of antibiotics against drug resistant pathogens that also disrupt the preformed biofilms, making the treatment vastly effective and efficient.

To the best of our knowledge, this study for the first time characterized and evaluated antibacterial and anti-biofilm activities of previously identified new compounds by our group (Mohapatra et al., manuscript communicated) against susceptible as well as clinical isolates of Gram-negative pathogens.

## CONCLUSION

The results showed that new compounds produced by SJ33 strain reported in our previous study possesses antibacterial and anti-biofilm potential against Gram-negative bacteria such as *K. pneumonia* and *E. coli*, which are prominent biofilm formers. The leakage of cellular contents and membrane disruption indicated bactericidal efficacy of compounds against *K. pneumoniae*. In addition, the compounds C1 and C2 dispersed established mature biofilms of *K. pneumoniae* MTCC109 as well as clinical isolate of *K. pneumoniae* at sub-MIC. Recently, it has been highlighted that antibacterial agents are better candidates for treatment of chronic infections caused by biofilm forming pathogens. In conclusion, antibacterial compounds C1 and C2 have demonstrated significant anti-biofilm activity against susceptible as well as drug resistant pathogens and can also be used for the development of novel anti-infective therapeutics in future. Further detailed mechanistic studies are required on these reported low molecular weight compounds to fully understand their nature and beneficial potential in various clinical applications.

## Acknowledgements

Authors acknowledge DST-FIST and UGC-SAP for the infra-structure facilities through special grants to the Department of Biochemistry & Molecular Biology, Pondicherry University, India. We are also thankful to IIT Madras, Chennai, India for SEM analysis. Authors are grateful to University Grant Commission (UGC), New Delhi for the financial support to ARM.

## Supplementary Table

**Table S1. Inhibitory Spectrum of antibacterial compounds C1 and C2 against both susceptible and multi-resistant Gram-negative pathogens.** Inhibition zone in millimetres (mm) inclusive of well diameter 6 mm; values are shown in mean ± SD of three independent experiments performed in triplicates.

