## Supplementary Information for "Anti-biofilm activity of new low molecular weight compounds produced by *Lactiplantibacillus plantarum* SJ33 against *Klebsiella pneumoniae*"

School of Life Sciences

Pondicherry University,

R. V. Nagar, Kalapet,

Puducherry – 605014,

**Table S1. Inhibitory spectrum of antibacterial compounds C1 and C2 against both susceptible and drug resistant Gram-negative pathogens.**

| **Indicator Bacterial Strains** | **Zone of Inhibition** | |
| --- | --- | --- |
|  | **C1** | **C2** |
| ***K. pneumoniae* MTCC 3384** | 15±1 | 14±1 |
| ***K. pneumoniae* MTCC 109** | 16±1.5 | 18±2.3 |
| **Resistant *K. pneumoniae*** | 15±0.5 | 14±0.6 |
| ***E. coli* MTCC 728** | 17±1.1 | 15±0.6 |
| **Resistant *E. coli*** | 17±2 | 16±4.3 |

Inhibition zone in millimetres (mm) inclusive of well diameter 6 mm; values are shown in mean ± SD of three independent experiments.

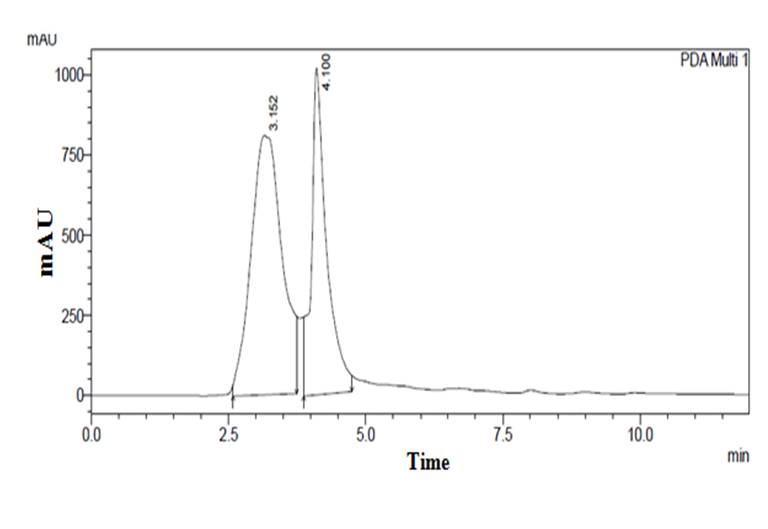

**Fig. S1 Analytical reverse phase HPLC profile monitored at 254 nm and structure of compounds (a) C1 and (b) C2.**

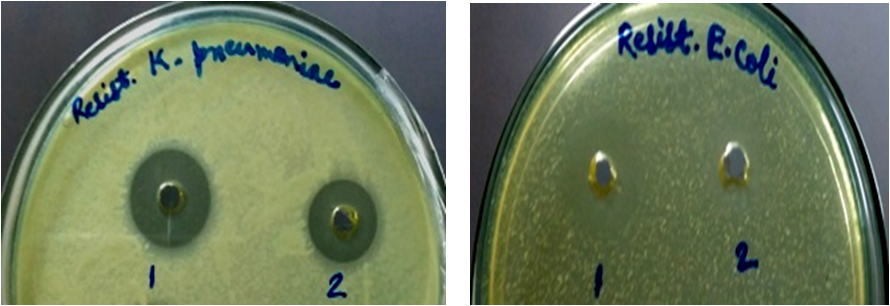

**Fig. S2 Antibacterial activity of identified compounds against clinical isolates of *K. pneumoniae* and *E. coli*.** Here, 1 indicates compound C1 and 2 indicates compound C2.

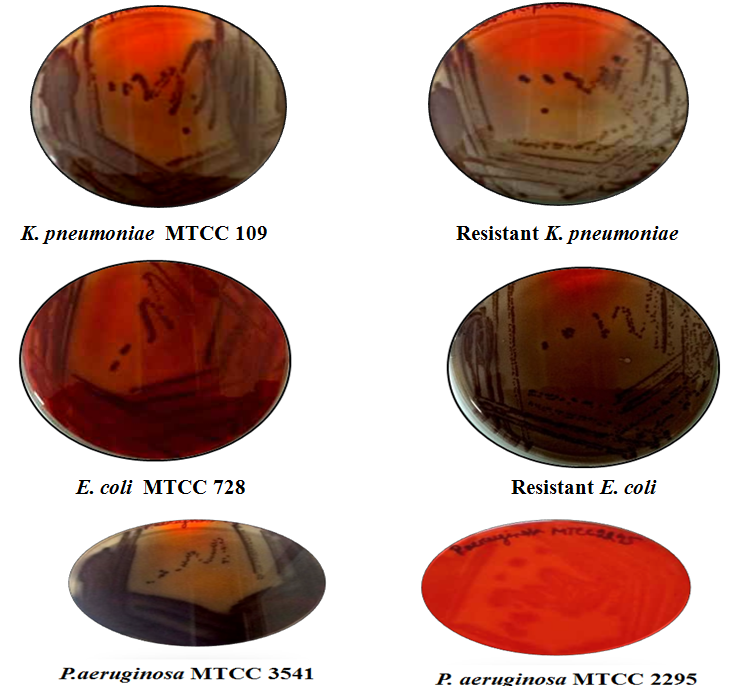

**Fig. S3 Exopolysaccharide production by susceptible Gram-negative bacteria as well as clinical isolates of *K. pneumoniae* and *E. coli* using congo red assay.** Black colonies on congo red agar shows biofilm formers, whereas white or pink colonies indicates non biofilm forming pathogens.
